# Silicon as a microfluidic material for imaging and incubation of droplets

**DOI:** 10.1101/2022.09.09.507341

**Authors:** Nicolas Lobato-Dauzier, Robin Deteix, Guillaume Gines, Alexandre Baccouche, Benediktus Nixon Hapsianto, Shu Okumura, Guilhem Mariette, Djaffar Belharet, Samuel Queste, Laurent Jalabert, Matthieu Denoual, Yannick Rondelez, Hiroshi Toshiyoshi, Hiroyuki Fujita, Soo Hyeon Kim, Teruo Fujii, Anthony J. Genot

**Affiliations:** LIMMS, CNRS-Institute of Industrial Science, IRL 2820, University of Tokyo, Tokyo 153-8505, Japan; Institute of Industrial Science, The University of Tokyo, Meguro, Tokyo 153-8505, Japan; Laboratoire Gulliver, CNRS UMR 7083, ESPCI Paris, PSL Research University, Paris 75005, France; Institut FEMTO-ST, Université Bourgogne Franche-Comté, CNRS UMR 6174, Besançon 25030, France; GREYC, CNRS UMR 6072, ENSICAEN, Caen 14000, France; Advanced Research Laboratories, Tokyo City University, Setagaya, Tokyo 158-0082, Japan

**Keywords:** Droplet microfluidics, droplet chamber, Silicon, enzymology

## Abstract

Droplet microfluidics has become a powerful tool in life sciences, underlying digital assays, single-cell sequencing or directed evolution, and it is making foray in physical sciences as well. Imaging and incubation of droplets are crucial, yet they are encumbered by the poor optical, thermal and mechanical properties of PDMS - the *de facto* material for microfluidics. Here we show that silicon is an ideal material for droplet chambers. Si chambers pack droplets in a crystalline and immobile monolayer, are immune to evaporation or sagging, boost the number of collected photons, and tightly control the temperature field sensed by droplets. We use the mechanical and optical benefits of Si chambers to image ∼1 million of droplets from a multiplexed digital assay - with an acquisition rate similar to the best in-line methods. Lastly, we demonstrate their applicability with a demanding assay that maps the thermal dependence of Michaelis-Menten constants with an array of ∼150,000. The design of the Si chambers is streamlined to avoid complicated fabrication and improve reproducibility, which makes Silicon a complementary material to PDMS in the toolbox of droplet microfluidics.

**Significance Statement:** As the technological engine behind single-cell sequencing and digital assays, droplets microfluidics has revolutionized life science and molecular diagnosis, and is making foray into physical sciences as well. Observing droplets in a controlled manner is becoming crucial, but PDMS - the *de facto* material of microfluidics – hampers imaging and incubation. Here we revisit silicon as a microfluidic material and show that its superior mechanical, optical and thermal performances improve the throughput and operation of droplets assay.

## Introduction

In the past decade, emulsions of droplets have enabled groundbreaking advances in life sciences. Their massive generation rate (∼100Hz-10MHz^1,2^), their monodispersity (a few %) and their minute volume (∼1-1000 pL) make droplets ideal reactors for applications that need high-throughput, low consumption of reagents and quantitativeness. Droplets microfluidics has become routine in single-cell analysis^3–6^, digital PCR^7–9^ or directed evolutions ^10–14^, where it is used to digitally encapsulate oligonucleotides, plasmids or cells. Droplets are also finding applications in other fields, ranging from chemical synthesis^2,15,16^ and nonlinear chemistry^17–20^, enzymology^14^, drug screening^21^ and toxicology^22^, to microbiology^23^, cell biology, and tissue engineering^24^. In recent years, the use of droplets has even extended to study the physics of crystallization, phase separation, gelation and colloidal aggregation^25–27^.

In typical end-point assays (e.g. digital assays), droplets are first generated in a PDMS chip, incubated in a controlled temperature profile off-chip (e.g. in thermocycler), and then interrogated with a dedicated device. In other applications, timing is important and droplets must be continuously imaged while incubated (time lapse imaging) - for instance to monitor cellular growth or gene expression ^28^. Lastly, some advanced applications call for both temporal *and* thermal resolution - for instance in enzymology, directed evolution, phase separation, or colloidal self-assembly - where kinetics and temperature are tightly linked.

However, it remains challenging to continuously image a large population of droplets, even more so when their temperature must be controlled. In-line methods (e.g. droplets cytometry^29,7,30,31^) circulate droplets inside a channel and read their fluorescence with detectors located at fixed positions in the channel. In-line methods offer a high throughput but with few time points and over a short duration (being limited by the residency time of droplets in the channel). They also lack morphological information about the content of the droplets, are difficult to set up and multiplex in several colors, and are hardly compatible with temperature-resolved measurements where each droplet experiences a different temperature. On the other hand, wide-field imaging works the opposite way: droplets are immobilized in a chamber and scanned by a moving microscope stage. Wide-field imaging brings significant advantages. First, it is easy to set up and automatize: commercial fluorescence microscopes can automatically scan a droplet array in several colours with limited hands-on time. Secondly, it is thermally, spatially, and temporally resolved: each droplet can be kept at a constant and distinct temperature (when placed in a temperature gradient), and repeatedly imaged with a spatial resolution of ∼1-10 μm and a time resolution ranging from milliseconds to minutes or hours. Lastly, wide-field imaging also offers a high-throughput thanks to the large surface of CMOS cameras, which capture tens of thousands of droplets in a single field of view.

The microfluidics community has designed a variety of droplet chambers to maximize the benefit of wide-field imaging. While some chambers have been realized in materials like glass^18^ or PMMA^32^, most have been made from PDMS, the standard material for prototyping in academic microfluidic. Starting with the DropSpot array in 2009, droplet chambers made of PDMS have been reported for application in digital PCR, live-cell imaging, screening of biomolecules, molecular diagnostics or microbiology ^28,33–40,24,41–44^.

While these PDMS chambers enabled groundbreaking studies, they kept running into the same limitations of PDMS. First, PDMS is not mechanically rigid, which makes it difficult to fabricate large freestanding chambers because they collapse under their own weights. This was addressed by adding supporting structures like pillars^34,45^, but at the cost of reducing the filling factor, which often falls short of the maximal packing density of droplets (∼90% in 2D). Secondly, the poor thermal conduction of PDMS makes it delicate to control the temperature field sensed by the droplets. Thirdly, PDMS expands upon heating and is not airtight, which causes evaporation and displacements of droplets during incubation. Lastly, the autofluorescence of PDMS degrades the optical signal, especially when the droplets must be imaged through the PDMS slab.

The ideal droplet chamber should meet stringent mechanical, thermal, optical and storage criteria. The chamber should maximize the packing density of droplets. It should be mechanically robust and airtight to prevent the compression, stacking, displacement or evaporation of droplets, and should not thermally expand when heated. The chamber should be wide enough to store millions of droplets across its surface, but be sufficiently thin and flat to hold only a single droplet across its depth - which imposes an extreme form factor and requires mechanical rigidity. The ideal chamber should be thermally conductive to keep the temperature locally uniform and quickly transmit temperature jumps to the droplets (like in PCR cycling). Lastly the chamber should not autofluoresce.

We reasoned that Si - a crystalline material - would make better droplet chambers than PDMS - an elastomer - because it is more mechanically rigid, more thermally conductive, more reflective and less permeable to water vapor (Table S1). The surface of silicon is flat, and easy to micro pattern and modify, for instance to tune wettability^46,47^. Those benefits have led Si to be used as substrate for industrial DNA synthesis ^48^. Digital PCR in arrays of Silicon microchambers (where the PCR reaction is directly done in a micrometric Si chamber rather than in droplets) has been demonstrated – highlighting the good thermal performance of Silicon^49^.

Yet PDMS has largely been preferred to Si for microfluidic devices due to the (perceived) complexity of silicon processing. Firstly, silicon chips must be processed in a cleanroom and cannot be replicated from a master mold like PDMS chips. Secondly, bonding silicon chips to glass (*e*.*g*. anodic bonding) and connecting them to the outside world (by drilling or back-etching access ports) is undeniably more complex and expensive than for PDMS chips (which just needs punching and tubing). While those points apply to classical microfluidic silicon devices (e.g. for generating droplets^2^), we reasoned that they would not necessarily apply to silicon devices for storing droplets. If the design of Si chambers could be streamlined to remove complicated processes (bonding, drilling and back etching), then silicon could become a competitive alternative to PDMS. Here we report silicon droplet chambers for the imaging and incubation of droplets. The silicon chambers enjoy superb mechanical, optical and thermal properties - making them ideal for imaging digital assays and time-resolved thermal studies of biosystems.

## Results

### Fabrication and filling

The design of the chamber is shown in Fig. 1. A chamber is made of a square or rectangular recess in a silicon wafer, surrounded by short inlet and outlet channels for filling, and covered by a coverslip. The depth of the chamber matches the diameter of the droplets - forcing them to spread into a monolayer. The silicon block is noticeably thinner (∼0.5-1mm) than a PDMS slab (∼1-5 cm), while being more thermally conductive and mechanically rigid.

**Figure 1:**
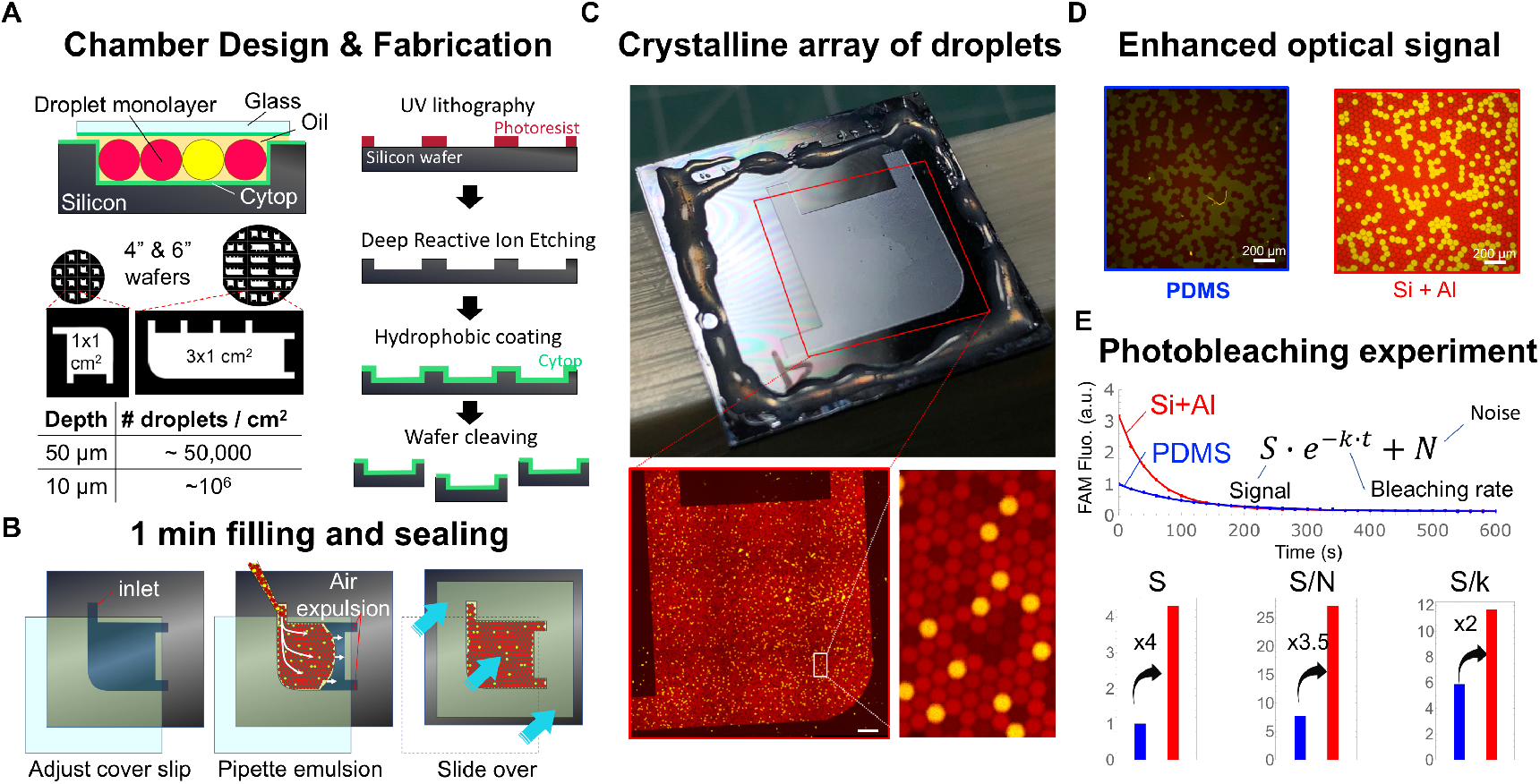
Fabrication, filling and optical observation of silicon droplets chambers. **A**, Fabrication. The chambers are fabricated by standard processing in the clean room. The depth of the chamber is tightly controlled by the etching time. After fabrication, the chambers are cut out by breaking the wafer along its crystal planes, or by slicing the wafer. **B**, Filling (Extended Video 1). The chamber is covered with a coverslip, and filled from one of the inlets, forcing air out of the chamber. After filling the chamber is closed either by capillary forces, or with epoxy glue **C**, Chamber after filling and sealing with glue (Extended Image 1). The chamber contains ∼50,000 droplets that are organized in macroscopic crystalline domains. **D**, Optical enhancement. An emulsion of droplets containing Rhodamine, alexa 647 and FITC dextrans was imaged under identical conditions in a Si/Al chamber and a PDMS chamber for reference. **E**, Prolonged illumination bleaches FITC, yielding the optical background (N), the bleaching rate (*k)*, and the optical signal (S). From this, we extract the signal-to-noise ratio (S/N), as well as the total number of photons collected (S/k, expressed in arbitrary units).

The silicon chambers were fabricated with a streamlined process: spin-coating of the resist, UV exposure, resist development, deep reactive ion etching, and wafer slicing (and those steps can be automated in a modern cleanroom). Since the chamber does not contain micrometric features in the xy plane, a chrome mask was unnecessary and we used an inexpensive plastic film as a mask for prototyping. Complex processing was not needed either (e.g. mask alignment, anodic bonding, back-etching or drilling). Overall, we fit ∼30 chambers of 1 cm^2^ on a 6” inch wafer, which brings the fabrication cost of a single chamber to ∼5-10€ in an academic cleanroom (including wafer cost and processing time).

The depth of the chamber - controlled by the etching time - is highly uniform despite the extreme form factor of the chamber (∼200-1000). We profiled the depth of a dozen of chambers distributed along the diameter of a 6-inch wafer (Table S2). For a nominal depth of 50 μm, the measured depth at the center of each chamber is close to the nominal depth, and varies little between chambers (49.6 +/-2.1 μm). Inside a chamber, the depth displays a slightly concave profile with a trenching on the edges. The depth deviates by ∼3 μm on average between the center and the edge of the chamber, which represents ∼6% of the nominal depth (50 μm) and 0.03% of the length of the chamber (1 cm). (Results are similar for the 10 μm chambers, Table S2).

The uniformity of depth and the mechanical rigidity of silicon ensures that droplets do not form a bilayer during filling, and that they do not get compressed or dislodged during incubation. It would be almost impossible to get the same uniformity and mechanical rigidity from a free-standing PDMS chamber. PDMS is an elastomer that notoriously sags under its own weight, which makes the fabrication of PDMS structures with a large aspect ratio difficult ^50^. Even with supporting structures like pillars to prevent the deformation or collapse of the chamber, the depth profile of a PDMS chamber is still determined by the thickness profile of the SU8 master, which is hard to uniformize because it is governed by spin-coating, and problems like edge beads, striations, dusts, and air bubbles are common occurrence in spin-coating that impact the uniformity of thickness.

### Filling and sealing

Filling and sealing the chamber are easy. Filling does not need specialized equipment (such as a pressure pump), but rather relies on physical forces that pack the emulsion into a crystalline monolayer of droplets. First, a coverslip is laid on top of the chamber (both coated with a hydrophobic polymer), leaving the inlet and outlets open. The water-in-oil droplet emulsion is then pipetted in the chamber through the inlet. The emulsion fills the chamber by capillarity, expelling air through the outlets. The corner opposite the inlet and outlets is rounded to prevent air bubbles from stagnating. After filling, the coverslip is gently slid over the silicon block to cover and close the inlet and outlets (this sliding is facilitated by the flatness of silicon). The chamber is then sealed, either by capillarity (by leaving a film of oil around the chamber), or by gluing the edges of the coverslips to the silicon bloc. Capillary closing is sufficient for incubating the array at moderate temperatures (<60°C), and allows for retrieval of the droplet arrays. Glue improves the air tightness and mechanical stability when chambers are heated at high temperatures (∼60-80°C, vide supra), but makes retrieval more involved.

Although filling is simple, it is remarkably efficient. The droplets self-organize into a crystalline monolayer whose large domains can be directly seen with the naked eye (Fig. 1C). Under the microscope, the droplets are packed into a honeycomb lattice, an arrangement which maximizes storage density. There are few visible air bubbles (a common issue in droplets microfluidics), and the droplet arrays remain hydrated at room temperature for days or weeks, without signs of coalescence. The simplicity of filling and sealing contrasts with the usual practice for connecting silicon chips to the real world, which is to etch or drill the back of the wafer (to open access ports for filling), and to seal the device with anodic bonding^51^. Back etching is more time-consuming than front etching because the whole wafer must be etched through (∼0.5-1 mm vs 50 μm). In addition, anodic bonding requires clean and flat surfaces (in addition to high temperatures and large voltages), which complicates fabrication and increases the likelihood of failed devices. Tubing is not necessary either, which further simplifies operation.

### Optical enhancement

While the Si chamber blocks transmitted light, their superb optical properties make them ideal for epifluorescence imaging - the readout modality for the vast majority of droplet assays. Note that if brightfield imaging is needed, it would need to be conducted in reflected mode rather than transmission mode.

First, Si chambers are mechanically rigid and can be sealed with a thin coverslip of standard thickness (∼170 μm) - making the chambers compatible with the vast majority of objectives (Figure S1). By contrast, PDMS devices are not rigid and are usually sealed with a thick (∼1 mm) glass slide for bonding and handling. But this millimetric thickness causes optical aberrations (most objectives correct aberrations for a thickness ∼170 μm) and imposes a long working distance - excluding lenses with a high numerical aperture and reducing resolution and brightness (Figure S1). However, a good numerical aperture is necessary to resolve the content of droplets, for instance to monitor inhomogeneous processes, like the polymerization, jellification or phase separation. This kind of measurements need to spatially resolve the content of the droplet on the microscale, which mandates a short working distance and high numerical aperture.

In addition, we hypothesized that the flat and reflective surface of the Si chamber enhances the optical signal by acting as a mirror^52^. The reflectivity of silicon is ∼40% (in the 400-600 nm wavelength range), and increases to almost 100% with the coating of aluminum (a simple clean room process that can be performed on several wafers at once). Assuming a perfect reflection from the surface, the flux of excitation photons coming from the objective and exciting the droplets is doubled, and the flux of photons emitted by the droplets and collected by the objective is also doubled, resulting in 4-fold enhancement of brightness. Noting that in epifluorescence microscopy the brightness of the sample scales typically like the 4th power of the numerical aperture^53^, this 4-fold enhancement is equivalent to increasing the numerical aperture of the objective by 40%. (e.g. collecting the light of a hypothetical 40X NA=0.55 objective while benefitting from the field of view of a 20X NA=0.4 objective).

We tested these assumptions by comparing the optical properties of a PDMS chamber and an Al-coated silicon chamber. We prepared an emulsion of identical droplets containing Dextran molecules conjugated to FITC fluorophores (a dye with fast photobleaching). After spreading the droplets in their respective chambers, we illuminated and imaged each chamber under identical conditions (Fig. 1D). As expected, the Al/Si chamber quadruples the photon flux *S* (as measured by the initial average fluorescence), and doubles the bleaching rate *k*, which overall doubles the number of photons S/k collected from the dyes over their lifetime. We were concerned that the fluorescence background could have been large because the excitation light is reflected back to the objective by the silicon surface. While this light is filtered out by the dichroic mirror and the emission filter, even a small leak could result in an overwhelming background. However, this turns out not to be a cause for concern, and the background *N* of the Si/Al chamber is only slightly worse (∼20%) than in PDMS chambers. Overall, the Si/Al chambers has an optical signal-to-noise ratio ∼3 fold larger than for PDMS chambers. This optical enhancement shortens the acquisition time and reduces illumination power, allowing to scan more droplets in a given time and to reduce phototoxicity. It also boost the apparent numerical aperture of the objective, which improves resolution^52^. This could improve the imaging of fine structures inside droplets, for instance to distinguish the morphology of cells or colloidal aggregates^27^.

### Imaging of a digital assays

We then exploited the mechanical rigidity and optical enhancement of the Si chamber to quickly image a large array of droplets from a digital assay, one of the major applications of droplet microfluidics^8,9,34,54^. The turnaround time, limit of detection and dynamic range of a digital assay are directly related to the size and number of droplets imaged. Small droplets (∼10 μm) yield quicker reactions (by raising the effective concentration of a single molecule) and digitalization through Poisson encapsulation, while large populations of droplets (∼10^4^-10^5^) give a wider dynamic range and a lower limit of detection.

We used a Si chamber to image a multiplexed population of droplets (∼1 million of 10 μm droplets), and capture in a single run the dilution curve of an isothermal digital assay for microRNA quantification^54^. The whole chamber with ∼900,000 droplets was imaged in 4 colours in about ∼25 minutes - an average acquisition time of ∼420 μs/droplet/colour. Although acquisition was not optimized (most of the acquisition time was spent by the software on focusing), this throughput compares favorably with some of the fastest reported throughput in droplet cytometry (∼500 μs/droplet/colour^29,31^). The hands-on time (filling of the chamber and setting up of acquisition) was ∼10 minutes, which again compares favorably with droplet cytometry (which can take much longer to set up and run). However, we noted that filling the Si chamber with 10 μm droplets was slightly more difficult than with 50 μm droplets, which is likely due to the increased resistance in very thin layers of fluid.

### Thermal enhancement

We exploited the thermal properties of the Si chambers for high-throughput enzymology. In molecular diagnosis and enzymology, a deviation of a few degrees from the designed temperature can change the enzymatic activity and alter the validity of the results^55^. Yet droplets are often incubated in materials that are poor thermal conductors: mostly PMMA or Polypropylene (PP) when droplets are incubated off-chip in PCR tubes, or PDMS and glass when they are incubated on-chip.

Silicon chambers enjoy excellent thermal properties by virtue of their high thermal diffusivity (*D*∼90 mm^2^/s), which is on par with copper (∼111 mm^2^/s), ∼300 fold larger than glass (∼0.34 mm^2^/s), and ∼1000 larger than PDMS, PP or PMMA (∼0.10 mm^2^/s). In addition to being an excellent thermal conductor, Si has other advantages over PDMS for prolonged heating: it expands minimally when heated and has a low gas permeability (a common cause of evaporation in PDMS chambers).

A thermally diffusive chamber improves the incubation of droplets. First, it reduces the bias between the temperature set by the heater and the actual temperature felt by the droplets, which can reach ΔT=0.75°C for a glass slide (Supplementary Information). Secondly, it shortens the time for thermal equilibration time. The timescale *τ* to equilibrate temperature over a thickness of *L* is *τ*∼*L*^*2*^*/D*. For L=1 mm, *τ* ∼10 ms for Si, but on the order of *τ* ∼10 s for material like PDMS or polypropylene (a timescale that is matched by measurements of equilibration time in PCR tubes^56^). Lastly, a good thermal conductor maintains a uniform temperature over the droplet array by smoothing out local sources or sinks of temperature (air pockets between the heater and the chamber, air convection in the room, air bubbles in the array, dust, local heating by the objective light and else). According to the steady-state heat equation, the contribution of a heat source to the temperature field T is attenuated in the Fourier space by a factor of *Dk*^*2*^ (where *k* is the spatial frequency). So, Si attenuates spatial inhomogeneities on a length scale ∼30 times larger than PDMS.

In order to fully exploit the thermal benefit of Si, we set up a thermal platform^57^ (Fig.3A) - which we operated in two temperature regimes (uniform or gradient). In the first regime, the temperature is uniform in space (the same temperature is imposed to both Peltier elements), and either kept constant in time (for incubation) or varied over time between room temperature and ∼80 °C (*e*.*g*. to establish the melting curves of DNA strands). In the second regime, we established a stationary spatial gradient of temperature (by setting one Peltier element slightly above the room temperature and the other between 60 and 65 °C). In both cases, the temperature measured by the Pt100 sensors fluctuated by less than ∼0.015°C over the course of ∼10 minutes (Figure S2). We also mapped the spatial profile of temperature actually sensed by the droplets with *in situ* DNA thermometers^49,56^. We measure a linear temperature gradient of 4.9 °C/cm, close to the nominal gradient imposed by the Peltier elements (5 °C/cm). This confirms that thermal losses are negligible as silicon faithfully transmits the temperature field from the copper plate to the droplet array.

We tested the stability of an array of droplets to repeated temperature cycling. It is usually challenging to maintain the spatial integrity of a droplets array in PDMS upon heating because it is often accompanied by dislocation, merging or local evaporation of the droplets. This is due to the unfavorable properties of PDMS (thermal expansion and porosity) that cause movement and evaporation of the array. Evaporation in PDMS devices is usually addressed with fixes such as a vapor barrier (i.e., a glass slide embedded in the PDMS slab) or water tanks (hydrated channels running around the water chamber^28^), but those fixes complicate the fabrication and operation of the chamber.

In our heating experiment, we repeatedly cycled the Si chamber between 60°C and 80°C, and tracked the fiduciary droplets that register the local displacement of the droplet arrays (yellow in Fig. 3B). Satisfyingly, the vast majority of the droplets remain neatly packed and immobile during temperature cycling - in spite of the growth of a gas bubble near one of the outlets. The average displacement of fiduciary droplets is well below one droplet diameter (Fig. 3B), which we attribute to the mechanical rigidity of silicon as it is ∼1000 times less susceptible to thermal expansion than PDMS (Table S1). Silicon is also less permeable than PDMS to gas, and thus less susceptible to evaporation. The temperature stability of the chamber could be further improved by degassing the oil solution and emulsion before incubation.

### Enzymology

Lastly, we combine these properties (high-throughput and controlled temperature field) to measure *en masse* the temperature dependence of an enzymatic process - a fine-grained enzymology study which would have been difficult by other means. As a proof-of-principle, we measured the Michaelis-Menten curves for the enzymatic conversion step in our digital assay (Fig. 2A). In this step, a substrate (the input DNA strand) binds to a DNA template, which triggers its extension by a polymerase, and then nicking by nickase, releasing an output DNA strand. Although conversion is a two-step enzymatic process, it can be approximated as an apparent one-step process with the Michaelis-Menten equation. To map this dependence, we prepared with microfluidic scanning droplets with varying concentrations of the input DNA, keeping other reagents at a fixed concentration across the droplets^17,18^. We arrayed the droplets in a long Si chamber (3 cm x 1 cm, amounting to ∼150,000 droplets), placed them in a thermal gradient and imaged in time-lapse mode (Extended Video 2). After imaging, we binned the droplets by temperature, and for each bin we constructed the Michaelis-Menten curve relating the speed of the conversion reaction to the concentration of substrate. These curves yield for each temperature the apparent V_max_ (maximum production speed at full saturation of substrate) and K_M_ (concentration of input to reach V_max_/2). The velocity V_max_ is not monotonic with temperature and peaks around ∼ 47 °C.The K_M_ constant is well fitted by a Boltzmann law (Figure S3), suggesting that it is determined by a thermodynamically reversible process: the binding of the input DNA to the template. Those observation results would have been difficult with a low throughput process (such as bulk measurement in thermocycler), where only a few temperatures and only a few concentrations per temperature are tested.

**Figure 2:**
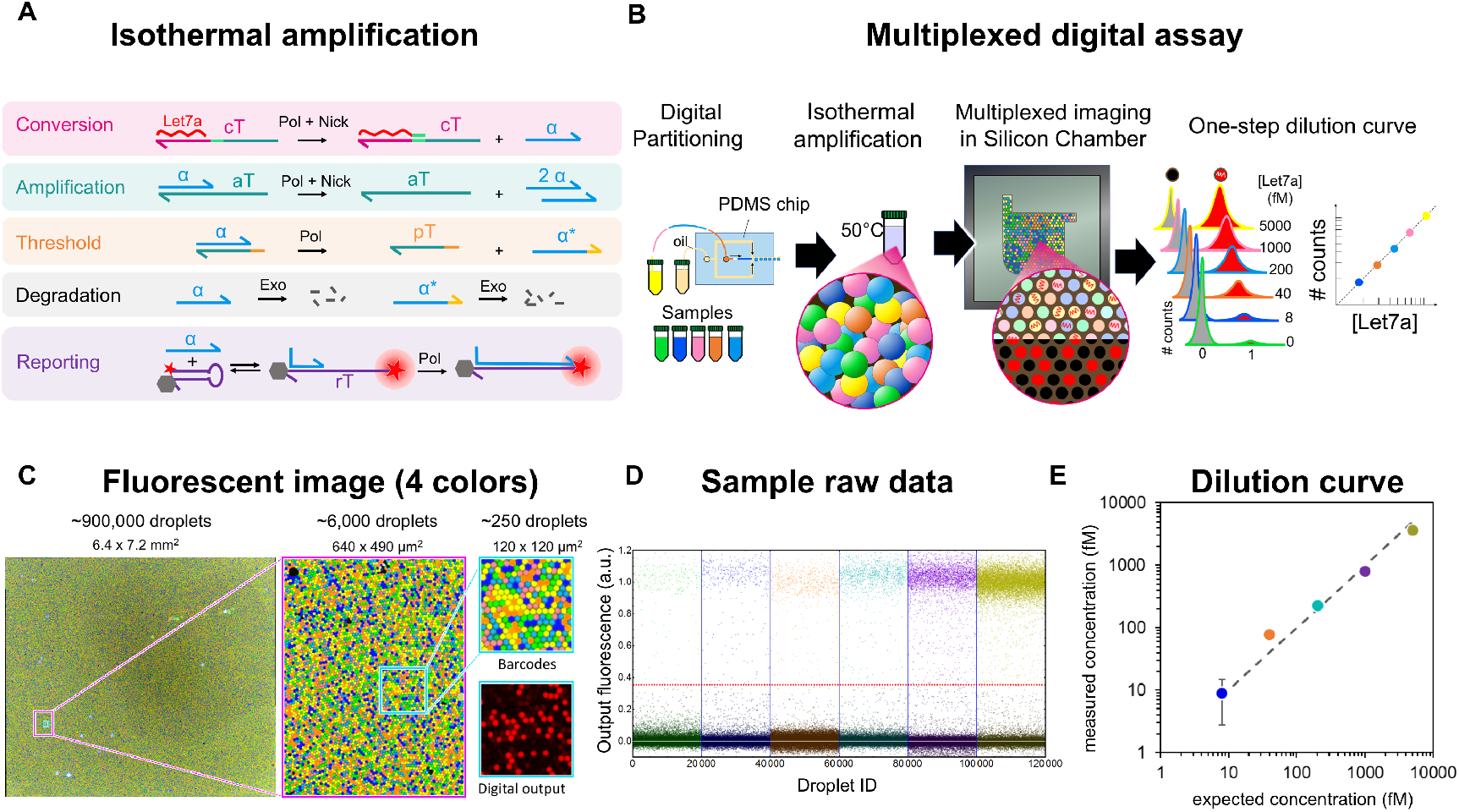
Imaging of a multiplexed digital assay with the Silicon chambers. **A**, Schematic of the isothermal assay. It comprises five molecular modules that exponentially amplify a target (*Let7a* miRNA) and report a molecular signal. **B**, Workflow. The isothermal assay is first digitally partitioned into droplets (*i*.*e* droplets contain either 0 or 1 strand of *Let7a*). We use a sample-changer to multiplex the assay and generate populations of droplets with varying dilution rates. After amplification, the droplets are arrayed in a Silicon chamber and imaged with multicolor fluorescence microscopy, which after processing yields the dilution curve of the digital assay. **C**, Raw image of the ∼1 million droplets array (Extended Image 2). Three colours encode the dilution rate of the droplet, and one colour reports the presence or absence of a Let7 strand. **D**, For each dilution rate, we plotted the output fluorescence of ∼20,000 randomly chosen droplets. **E**. Dilution curve plotting the measured concentration of Let7 against its expected concentration, for ∼500,000 droplets that passed quality filters. The error bars (only visible for the most diluted sample) show the standard deviation on the concentration measurement.

## Discussion

We have shown that silicon is an ideal microfluidic material for imaging and incubating droplets. The standardized fabrication process enhances reproducibility, as each chamber is made with precise and reproducible specifications. The chambers are simple to operate, and do not require specialized equipment. Lastly, they boast better optical, mechanical, thermal and storage performance than PDMS chambers. This makes silicon chambers ideal for experiments that mandate continuous imaging of many immobile droplets in a graded temperature field, for instance for high-throughput enzymology^58^.

We foresee various applications for the Si chambers in the life sciences: for real-time digital assays, directed evolution of enzymes, or live-cell imaging (thanks to their reduced phototoxicity, although long term incubation would be difficult due to the limited porosity to gases). The chambers could also benefit the physical sciences to map the phase diagram (temperature, concentration) of the many processes that can be followed with optical microscopy (colloidal self-assembly, phase separation, gelation, coacervate, solubility studies, crystallization). The chambers could also find application beyond droplet microfluidic, for instance in super-resolution microscopy where it could boost the number of photons collected and improve resolution for a modest cost and complexity.

## Materials and Methods

### Silicon chamber microfabrication

For the in-house process, 4 inch Si wafers (525 μm thick) were first cleaned with acetone, ethanol and water, dried and spin-coated with S1818 positive photoresist at 500 rpm for 30s followed by spinning at 3000 rpm for 30s and then baked at 110°C on a heat plate for 2 min. Photolithography was performed by directly adjusting a printed plastic (PET) film on top of the wafer to ensure that the chambers were aligned with the wafer orientation and exposing it to UV radiation (Union Optical, PEM-800 mask aligner) for 30s and developed for 1 min using a developer solution containing tetramethylammonium hydroxide (NMD-3, TMAH: 2.38 wt %). This rather long exposure and development help correct the imperfections from the PET mask. The wafer was then etched using Deep Reactive Ion Etching (DRIE) (SPT, Predeus Si deep reactive ion etching system). Chambers were etched to 110% of the desired droplet diameter to allow for a better flow of the droplets in the chamber while avoiding double layer. For the 50 μm chambers etching was performed by steps of 10 μm with 2 min break in between to prevent the wafer from overheating. Indeed, such overheating damages the photoresist resulting in particle deposition which after etching form pillars in the chamber. The Si wafer was finally cleaned with acetone, ethanol and water. and spin-coated with a hydrophobic fluoropolymer (CYTOP) at 500 rpm for 30s and 1000 rpm for 30s and baked at 180°C for 1h. Finally single chambers were obtained by cleaving the wafer.

To increase the fabrication throughput, we developed a process on 6 inch wafers (675μm thick) performed in a national academic cleanroom (FEMTO-ST, Besancon,France). The (100) face is the main plane for the fabrication process. The etching mask is S1813 photoresist 1.3μm thick deposited on a SussMicrotech ACS200. The dry etch is performed on an SPTS Rapier C2L etcher. We used different etching times depending on the desired etching depth. After etching, the photoresist was stripped with acetone and O_2_ plasma. The etching depths and homogeneities were characterized with a Dektak stylus profilometer. All the etching features with same dimensions are measured from bottom to top. Finally, all the structures are diced on a high precision dicing saw disco DAD3350.

### Samples preparation

Nucleic acid strands were purchased from Integrated DNA technology (IDT) or from Biomers and purified by high-performance liquid chromatography. All sequences are presented in Table S3-S6. Fluorescent Dextrans with a molecular weight of 10,000 Da were purchased from Thermo Fischer. Nb.BsmI, Vent (exo-), Bsm1, NBI enzymes as well as BSA were purchased from New England Biolabs (NEB). ttRecJ, a thermophilic exonuclease, was purified in the laboratory following a previously published protocol^59^. The working solution at 1.53 μM was obtained using Diluent A (NEB) and 1% Triton X-100, and stored at −20 °C. For each experiment, all common reagents were mixed into a mastermix to assure constant concentrations. We first added the DNA, RNA strands to the buffer and the Dextrans, BSA vortexed for 10s. For experiments using enzymes (Figure 2 & Figure 4), these were added last followed by a gentle vortexing and everything was assembled on ice to prevent an early start of the reaction. Once completed the mastermix was split into several tubes and we added the varying reagents as well as their respective fluorescent barcode. Bulk fluorescence (Figure 3C) was acquired using a CFX96 thermocycler (Biorad). The complete composition of each solution is bescribded in Table S7-S11

**Figure 3.**
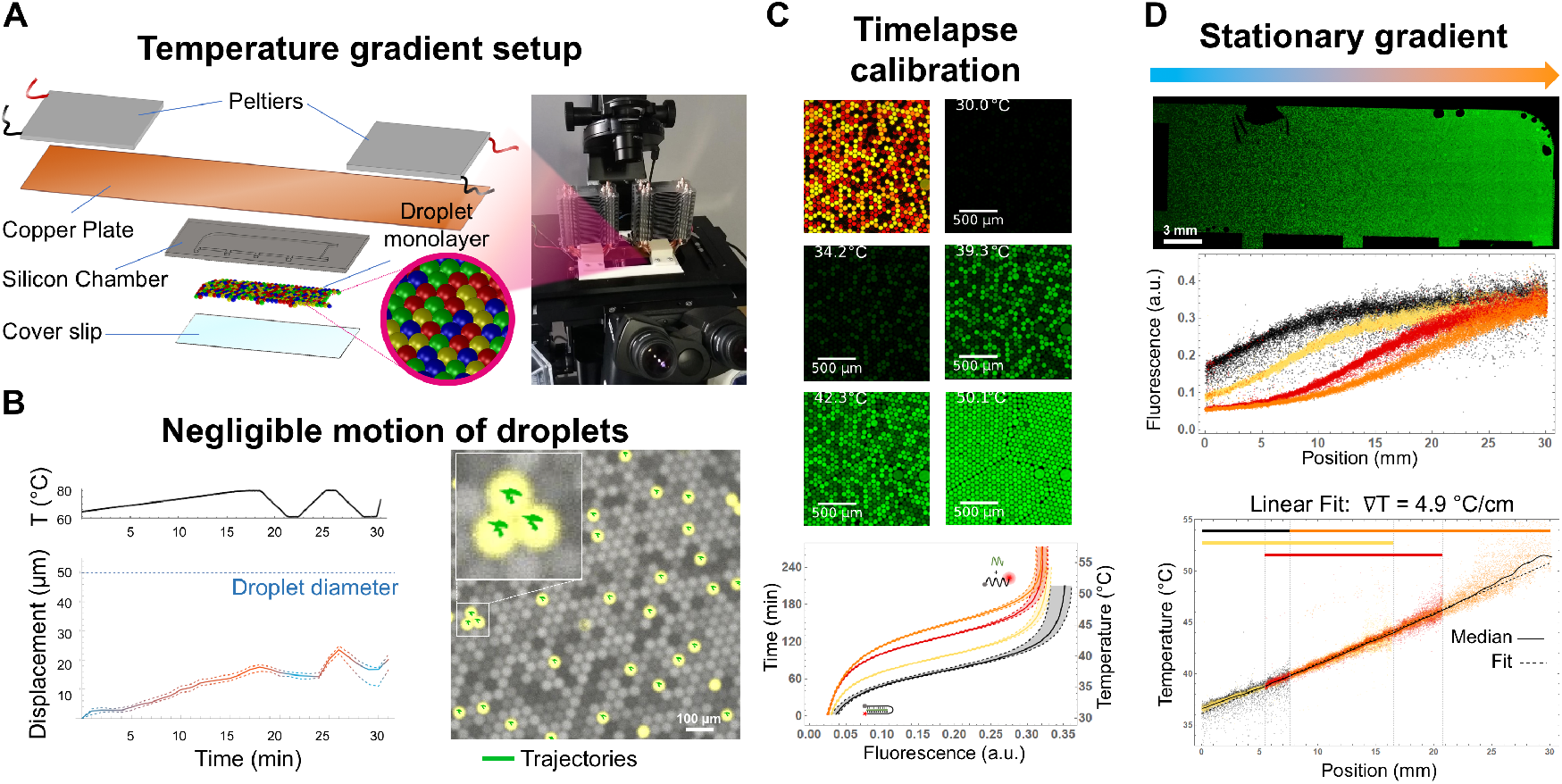
Thermal setup. **A,** Exploded view of the thermal setup. To gain control over the temperature field, we placed the Si chamber against a copper plate which is heated by two independent Peltier elements. The setup is encased in a 3D printed frame, and set inside the stage of the microscope. **B**, We measured the stability of the droplets array upon repeated temperature cycling between 60°C and 80°C (top plot). The yellow droplets are fiduciary droplets, and the green curves show their individual trajectories during heating and cooling (summarized in the bottom plot). **C**, Calibration of temperature measurements. We measured the temperature of droplets *in situ* with DNA thermometers, which are DNA nanostructures whose fluorescence responds nonlinearly to temperature. Each DNA thermometer maps temperature in a distinct range. We slowly heated an array of droplets with 4 distinct DNA thermometers and recorded the resulting fluorescence. **D**, In situ mapping of the temperature gradient. We placed an array of droplets with DNA thermometers in a stationary temperature gradient at 5°C/cm. Mapping fluorescence back to temperature reveals the temperature field in the droplets.

**Figure 4.**
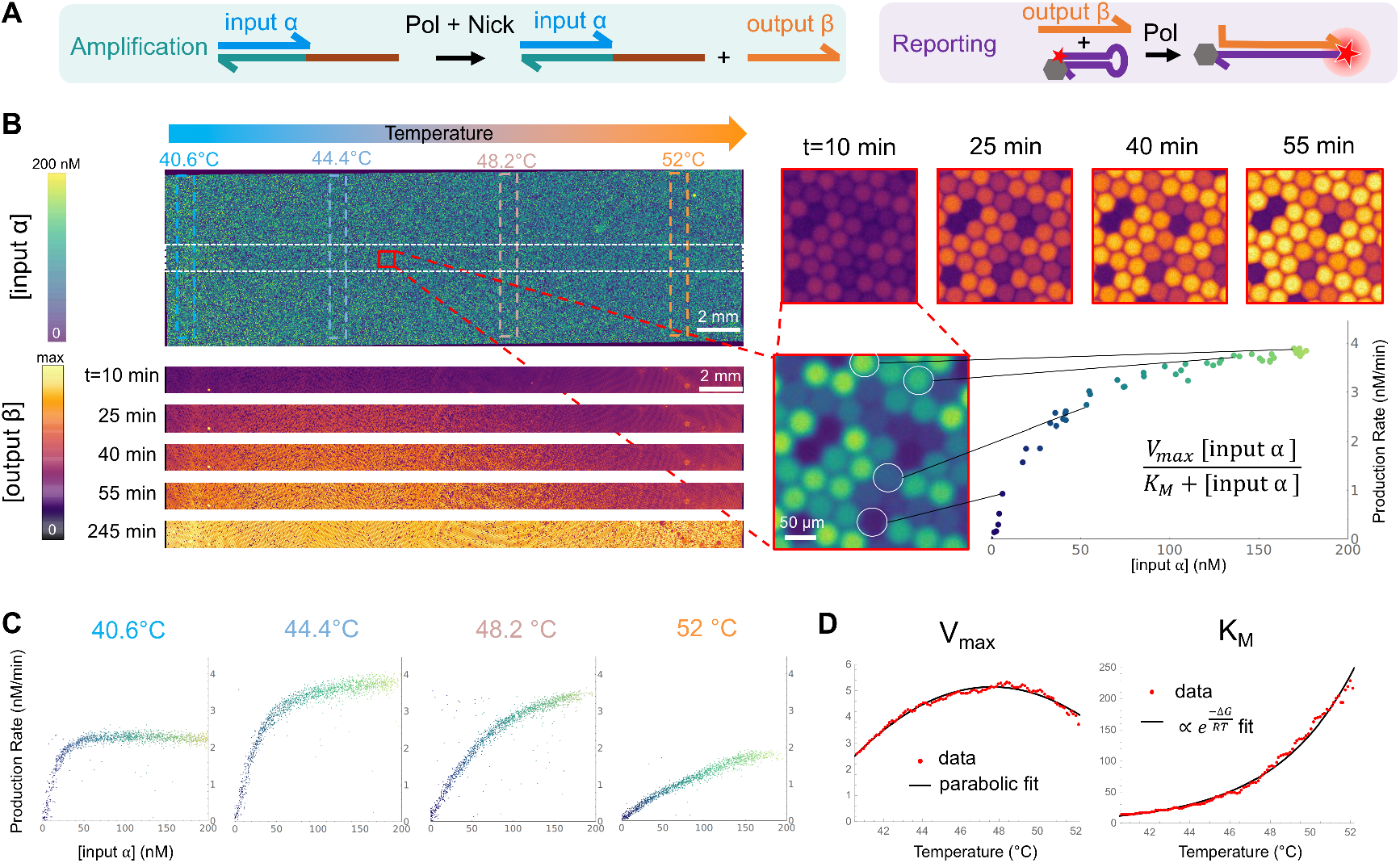
*En masse* thermal mapping of Michaelis-Menten constants. **A**, Enzymatic process studied. An input strand α (the substrate) binds to a DNA template, is elongated by a polymerase and nicked by nickase, releasing a strand (the product). The process is monitored by a separate reporter. **B**, Droplets with varying concentrations of α were prepared and arrayed in a Si chamber. The top image shows the array of droplets in the barcode channel (which is proportional to the concentrations of α). The bottom strips show the time evolution of the reporter’s fluorescence (Extended Video 2). The right side shows how a Michaelis-Menten curve is constructed. Each point corresponding to the derivative of the β fluorescence in a single droplet. **C**, Michaelis-Menten plots for various temperatures. The plot shows the production rate of β against the concentration of α. **D** Plots of Michaelis-Menten constant V_max_ and K_M_ for varying temperatures, extracted from the Michaelis-Menten plots (each data point in red corresponding to ∼1000 droplets).

### Microfluidic droplet encapsulation

We used an in-house microfluidic platform to generate water in oil droplets by hydrodynamic flow focusing using a pressure pump controller (MFCS-EZ, Fluigent, France) and PDMS microfluidic devices following previously published protocols: droplets of 50 μm diameter with discretely (Figure 1 & Figure 3) or continuously (Figure 4) varying content used a simple 3 channels PDMS device ^17,18^and 6 populations of droplets in the 10 μm range using a PDMS device with 2 different channel heights^54^ and a 3D printed sample changer^60^. Briefly, the PDMS devices were replicated using a silicon patterned with SU8 mold, and plasma bonded to a ∼1mm thick glass slide. They were then heated to 200°C for 5h to recover PDMS native hydrophobic state^61^. We used a fluorinated oil (HFE 7500, Novec) to which we added a surfactant (FluoSurf, Emulseo) at 3% w/v. The device is then placed onto a fluorescent microscope and each aqueous solution is plugged into a different inlet, all merging into one single channel that intersects with the oil channel. This produces monodisperse droplets whose size and composition can be finely tuned by changing the pressure ratio of water and oil. To create 50 μm droplets the pressure water:oil is 1:2. We scripted the pressure profiles to contribution of each aqueous channel, thus continuously varying the concentration inside droplets (Figure 4B). The droplets were collected in a pipette tip at the outlet before being transferred to the silicon chamber for imaging. This allows us to easily measure the volume of emulsion, collect separately different droplet populations or face any trouble without having to change device or to restart the whole experiment.

### Silicon chamber filling procedure

The collected emulsion is inserted in the silicon chamber(Figure 1B). A tutorial for the filling of the chamber is presented in Extended Video 1.

### Incubation and imaging

The silicon chamber was set under a copper plate (16 cm x 4 cm x 0.5cm) whose temperature is monitored using a Peltier controller (TEC-1122, Meerstetter) coupled to Pt100 sensors (RS-Pro, 10 mm x 2mm probe, 4-wire, Class A) and two Peltier elements (Adaptative, 40 × 40 mm ET-161-12-08-E) surmounted by a CPU cooler (Enermax, AM4 ETS–N31–02) as shown on Figure 3A. The chamber was imaged using a motorized Nikon Ti2-E epifluorescence microscope connected to an LED light source (pE-4000, CoolLED) and a sCMOS camera (Prime 95B 25 mm, Photometrics). We used 4x, 10x and 20x objectives (CFI Plan Apo Lambda NA: 0.2, 0.45, 0.75, Nikon) and filters corresponding to the desired wavelength (purchased from Semrock and Chroma). Large images were obtained by unshading with the BasiC plugin^62^ and stiching^63^ in ImageJ.

### Data analysis

Analysis from fluorescence images was performed using Mathematica as described in previous protocols^17,54^. Briefly, droplets were detected and tracked using the channel corresponding to a Dextran of constant concentration. Droplets composition were obtained by transforming barcodes fluorescence into concentration levels. Temperature was obtained using the position of each droplet and the value of the temperature gradient.

## Supporting information

SUPMAT for Silicon as a microfluidic material for imaging and incubation of droplets

Extended Image 1 Full chamber in Fig1C

Extended Image 2 Multiplexed Digital Assay Barcodes Image in Fig2C

Extended Video 1 Filling of Si Chamber

Extended Video 2 En Masse Thermal Mapping of Michaelis-Menten Constants

## Acknowledgments

We acknowledge support from the national Micro/nanofabrication network RENATER and the FEMTO-ST clean room in France as well as the VDEC cleanroom at the University of Tokyo. NLD, RD, SO acknowledges support from a MEXT studentship. AB acknowledges support form a JSPS postdoctoral fellowship.

