## Supplementary material for "Silicon as a microfluidic material for imaging and incubation of droplets": SUPMAT for Silicon as a microfluidic material for imaging and incubation of droplets

### Additional Figures and Tables

| Properties | PDMS | | Silicon | |
|---|---|---|---|---|
| Mechanical stiffness (Young modulus) (Pa) | $3.6\text{-}8.7 \cdot 10^{5}$ | [1] | $1.30 \cdot 10^{11}$ | [2] |
| Thermal conductivity ($W\ m^{-1}K^{-1}$) | 0.15 | [3] | 130 | [4] |
| Coefficient of thermal expansion α ($°K^{-1}$) | $200\text{-}300 \cdot 10^{-6}$ | [5] | $2.6 \cdot 10^{-6}$ | [6] |
| Reflectance (%) | 5-10 | [7] | 40-80 | [8] |

**Table S1 Comparison of some mechanical, thermal and optical properties of PDMS and silicon.**

| Chambers nominal depth | Measured depth at center | Measured depth at border | Measured depth max variation |
|---|---|---|---|
| 50 µm (N=15) | 49.5 ± 1.7 µm | 52.2 ± 2.3 µm | 3.5 ± 1.1 µm |
| 10 µm (N=15) | 9.6 ± 0.2 µm | 10.1 ± 0.3 µm | 0.6 ± 0.2 µm |

**Table S2 Characterization of the depth of the silicon chambers for the 6-inch wafer process.** Typical distance from center to border is 5 mm

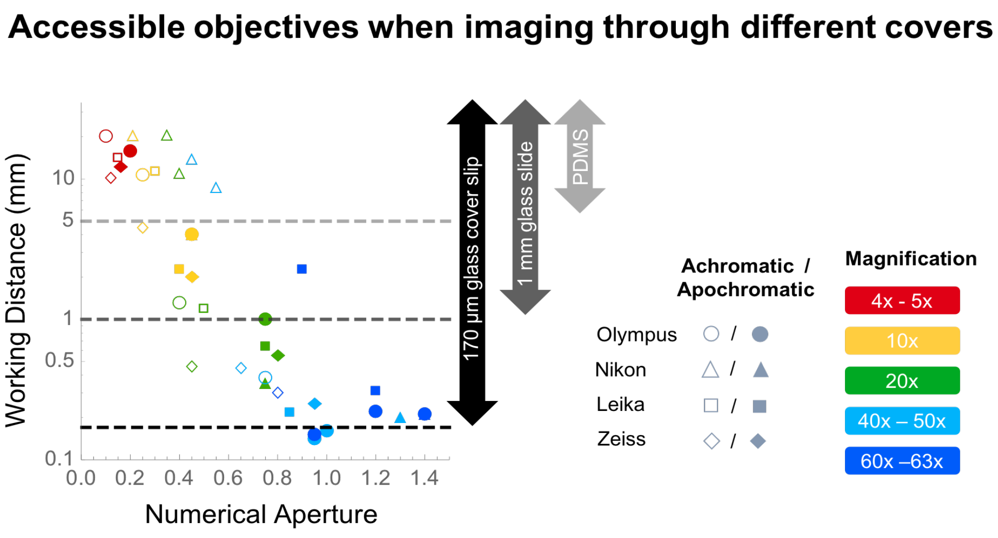


**Figure S1. Numerical Aperture and Working distance of typical microscope objectives.** The objectives with the best performances (Apochromatic with a large Numerical Aperture) are only accessible when using a 170 µm glass cover slip due to their shorter Working Distances. A slab of PDMS is typically a few millimeters thick, and the glass slides used to bond with PDMS are usually ~1 mm thick

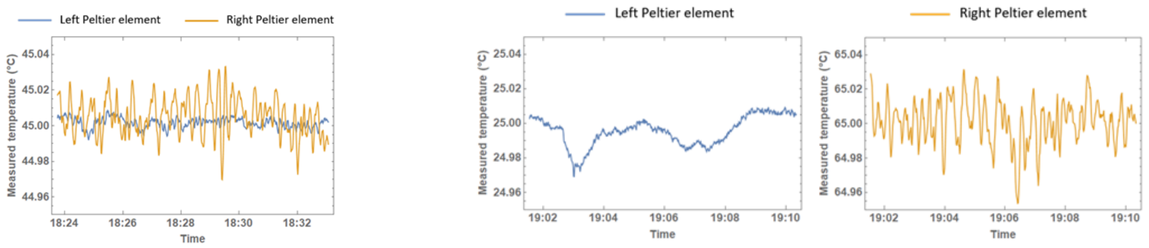


**Figure S2. Temperature stability of the Peltier setup in uniform (left) or gradient (right) mode.** The right Peltier element is noisier because being often used for heating 40°C above the room temperature which requires higher currents than on the left side.

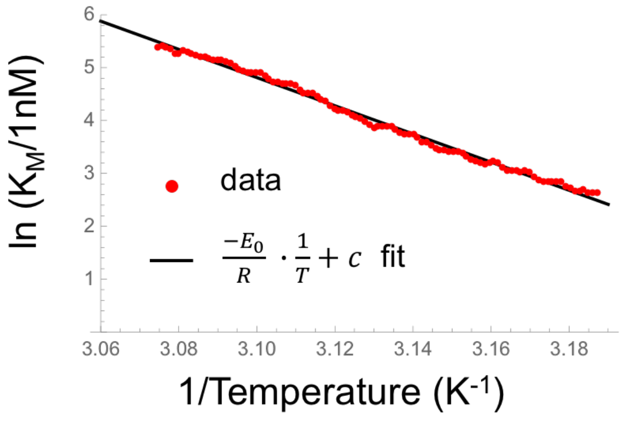


**Figure S3. Linear Fit of natural logarithm of $K_M$ against the inverse of the temperature.** This linear dependence shows that $K_M \propto e^{\frac{-E_0}{RT}}$ with $E_0 = 2.2 \cdot 10^5\ J$.

### Nucleic acid sequences

| Name | Sequence |
|---|---|
| Let-7a-RNA | UGA GGU AGU AGG UUG UAU AGU U |
| alpha: Bo12 | CATTCATCCCAG |
| aT: CBo12-2PS3 | C*T*G*GGATGAATGCTGGGATGAA |
| pT: pTBoT5S3P | T*T*T*TTCTGGGATGAATG |
| rT: rTBo-2BsmIAtto633 | Atto633*C*T*TCATGAATGCTGGGATGAAG BHQ2 |
| Let7atoBo-2+2P | TGCTGGGATGAAGTTTGACTCAAACTATACAACCTACTACCTCA |

**Table S3. DNA and RNA sequences for the multiplexed digital assay (Figure 2).** * denotes a phosphorothioate backbone

| Name | Sequence |
|---|---|
| Beacon Bottom: Cd11bex8 bottom | ATTACGAATTCACCAATGACGTAGCGAATGACTCCTAT-Atto647N |
| Beacon Top: Cd11bex8 top | BBQ 650-ATAGGAGTCATTCGCTACGTCATTGGTGAATTCGTAAT |

**Table S4. High temperature molecular beacon (Figure 3 B)**

| Name | Sequence |
|---|---|
| Clamp Switch | Atto 488-ATTTTCTTTTCCCCCCAGTTATTATTCCCCCCTTTTCTTTTG-BHQ1 |
| Stab 12 | AAAAGAAAAGGG |
| Stab 11 | AAAAGAAAAGG |
| Stab 10 | AAAAGAAAAG |
| Stab 9 | AAAAGAAAA |

**Table S5. DNA nanothermometers[9] sequences used in Figure 3 C,D**

| Name | Sequence |
|---|---|
| input α: X2 | CATTCACGATAG |
| output β: Ba12 | CATTCTGACGAG |
| cT: X2 to Ba12 | C*T*C*GTCAGAATGCTATCGTGAATG |
| rt: MB.Ba12 HEX | HEX-*T*T*CTGA TTTTCTCGTCAGAA-BHQ1 |

**Table S6. DNA sequences used for the thermal mapping of Michaelis-Menten constants.** * denotes a phosphorothioate backbone

### Solutions contents

| Optical enhancement & Photobleaching | |
|---|---|
| Component | Concentration |
| Additional mQ water | 70 % |
| NEB 3.1 buffer (10x) | 1x |
| dextran FITC | 100 nM |
| dextran Rhodamine | 0 / 100 nM |
| dextran alexa 647 | 100 / 0 nM |

| **1X NEB 3.1 buffer** (pH=7.9 @25°C) | |
|---|---|
| Component | Concentration |
| NaCl | 100 mM |
| Tris-HCl | 50 mM |
| MgCl2 | 10 mM |
| BSA | 100 µg/ml |

**Table S7. Composition of the droplets used for the optical characterization of the chamber (Figure 1 D,E)**

| Multiplexed digital assay | |
|---|---|
| Component | Concentration |
| mir Buffer | 25 µM |
| Let 7a | 0 / 8 / 40 / 200 / 1000 / 5000 fM |
| dextran Texas Red | 0 / 0 / 400 / 0 / 200 / 200 nM |
| dextran Cascade blue | 0 / 400 / 0 / 200 / 200 / 0 nM |
| dextran Alexa 488 | 400 / 0 / 0 / 200 / 0 / 200 nM |
| aT: CBo12-2PS4 | 50 nM |
| pT: pTBo12T5SP | 12 nM |
| cT: Let7atoBo-2+2P | 0.5 nM |
| rT: MB.Bo-2Bsm1Cy5 | 40 nM |
| BSA9000S | 200 µg/mL |
| Nb.BsmI | 300 u/mL |
| Vent(exo-) | 70 u/mL |
| BsmI | 7 u/mL |
| ttRecJ | 13 nM |
| ntBstNBI | 10 u/mL |
| in CFX96 @50°C | |

| mir Buffer | |
|---|---|
| Component | Concentration |
| NaCl | 100 mM |
| Tris-HCl | 50 nM |
| MgCl2 | 10 nM |
| BSA | 100 µg/ml |

**Table S8. Composition of the droplets used for the multiplexed isothermal assay (Figure 2)**

| Droplets tracking with heating | |
|---|---|
| Component | Concentration |
| Additional mQ water | 76% |
| Clamp buffer | 0.25 x |
| BSA9000S | 200 µg/mL |
| Beacon Top | 500 nM |
| Beacon Bottom | 500 nM |
| EvaGreen | 1x |
| Dextran Cascade Blue | 100 nM |
| Dextran Alexa 555 | 0 / 100 nM |

| Clamp buffer (pH=7) | |
|---|---|
| Component | Concentration |
| HEPES | 50 mM |
| NaCl | 300 mM |
| MgCl2 | 10 mM |

**Table S9. Composition of the droplets used for the characterization of the chamber at high temperatures (Figures 3 B & Figure 1 C)**

| Silicon chamber temperature calibration using DNA nanotermometers | |
|---|---|
| Component | Concentration |
| Additional miliQ water | 61 % |
| Clamp buffer | 1x |
| BSA9000S | 200 µg/mL |
| dextran Alexa 555 | 100 nM |
| Clamp Switch atto 488-BHQ | 100 nM |
| Stab 9 | 400 / 0 / 0 / 0 nM |
| Stab 10 | 0 / 400 / 0 / 0 nM |
| Stab 11 | 0 / 0 / 400 / 0 nM |
| Stab 12 | 0 / 0 / 0 / 400 nM |
| dextran Cascade Blue | 0 / 0 / 100 / 100 nM |
| dextran Alexa 647 | 0 / 100 / 0 / 100 nM |

**Table S10. Composition of the droplets used for the characterization of the thermal gradient Figure 3 C,D**

| *En masse* **thermal mapping of Michaelis-Menten constants** | |
|---|---|
| Component | Concentration |
| Additional miliQ water | 55% |
| Smix buffer | 1x |
| dNTP | 400 µM |
| rt: MB.Ba12 HEX | 200 nM |
| cT: X2toBa12 | 10 nM |
| dextran Cascade Blue | 200 nM |
| BSA9000S | 200 µg/mL |
| Nb.BsmI | 450 u/mL |
| Vent (exo-) | 70 u/mL |
| input α: X2 | 0 to 200 nM |
| Dextran Alexa 647 | 0 to 200 nM |

**Table S11. Composition of the droplets used for the mapping of Michaelis-Menten constants presented in Figure 4**

### Mathematical estimate of thermal loss

Let us consider a simplified model of heat losses: a material of thickness *L* has one of its sides in contact with a heater at temperature 60°C, and the other side in contact with air (with a temperature of 22°C far from the material). The material transfers heat to the environment (i.e. the air side) by convection and radiation. We assume that the material is thin and that the temperature of the side exposed to air is close to 60°C. The convective heat rate with air is given by Newton's law and we assume that the heat transfer coefficient is independent of the material and fixed by the surrounding air medium, for which we take a heat transfer coefficient of 10 W $m^{-2}$ $K^{-1}$ (typical for free convection). Then the material loses ~380 W $m^{-2}$ by convection to air. It also loses 224 W $m^{-2}$ by radiative transfer according to the Stefan–Boltzmann law (assuming an emissivity of 0.95). These heat losses are compensated by the establishment of a temperature gradient in the bulk of the material; the more thermally conductive the material is, the less the temperature drops. The differential of temperature ΔT between the hot and cold side of the material is $k/L\Delta T=P_{loss}$, where L is the thickness and k the thermal conduction. For glass, the temperature drop is ΔT=0.75 °C over a thickness of 1 mm, while the drop is negligible for silicon (less than 4 mK).
