## Supplementary figures and images for "Silicon as a microfluidic material for imaging and incubation of droplets"

### Extended Image 1 Full chamber in Fig1C

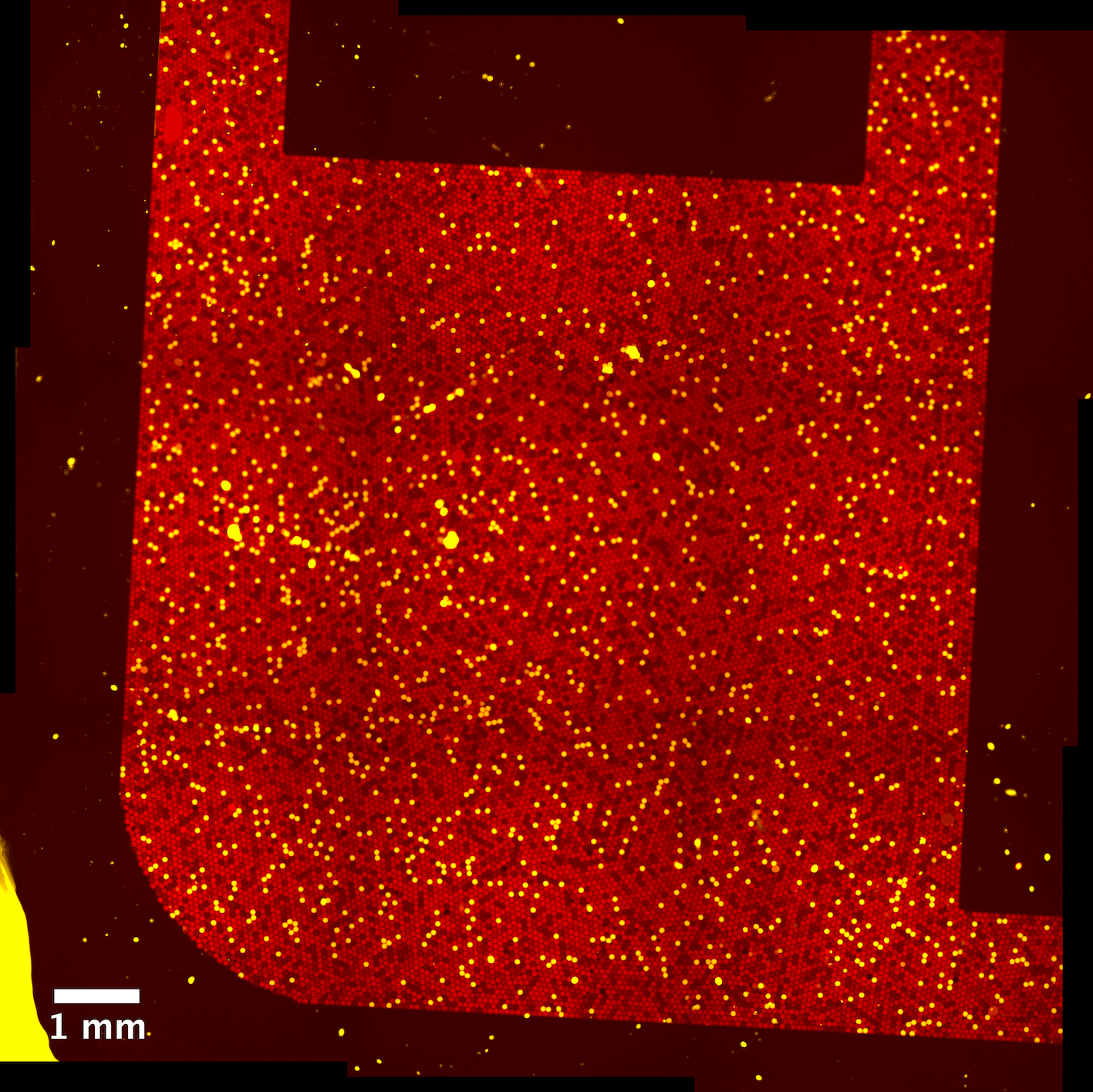

### Extended Image 2 Multiplexed Digital Assay Barcodes Image in Fig2C

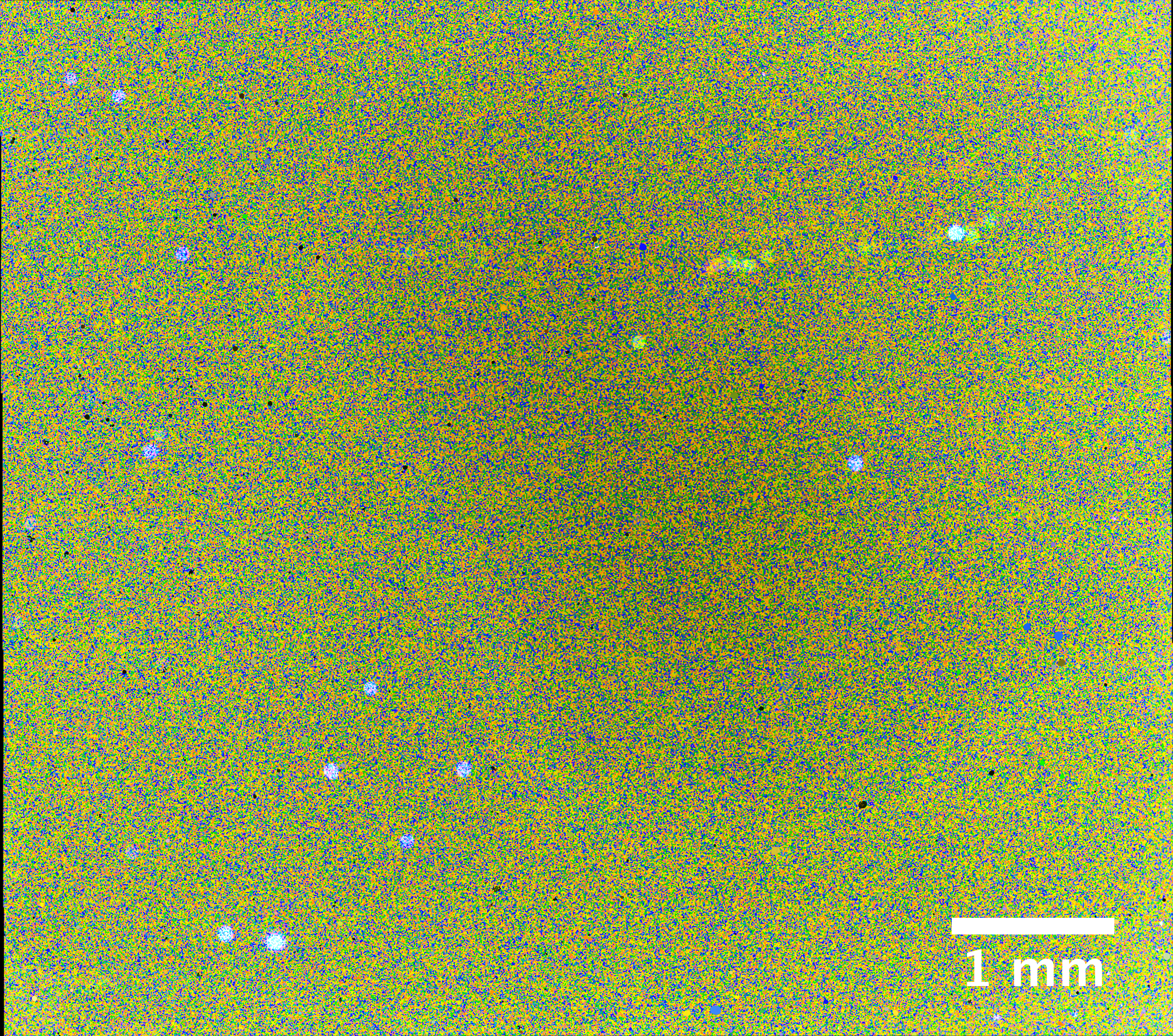
